# Massive analysis of 64’628 bacterial genomes to decipher a water reservoir and origin of mobile colistin resistance (*mcr*) gene variants: is there another role for this family of enzymes?

**DOI:** 10.1101/763474

**Authors:** Mariem Ben Khedher, Sophie Alexandra Baron, Toilhata Riziki, Raymond Ruimy, Seydina M. Diene, Jean-Marc Rolain

## Abstract

Since 2015, new worrying colistin resistance mechanism, mediated by *mcr*-1 gene has been reported worldwide along with eight newly described variants (*mcr*-2 to *mcr*-9) but their source(s) and reservoir(s) remain largely unexplored. Here, we conducted a massive bioinformatic analysis of 64’628 downloaded bacterial genomes to investigate the reservoir and origin of these *mcr* variants. We identified a total of 6’651 significant positive hits (aa sequence coverage > 90 % and similarity >50%) with the nine MCR variants from these genomes that include 39 bacterial genera and more than 1050 species. Although these variants could be identified in bacteria from human and animal sources, we found plenty MCR variants in unsuspected bacteria from environmental origin, especially from water sources. The ubiquitous presence of *mcr* variants in bacteria from water likely suggests another role in the biosphere of these enzymes as an unknown defense system against natural antimicrobial peptides and/or bacteriophage predation.

## Introduction

The global spread of multidrug-resistant bacteria (MDR) has greatly affected the effectiveness of antibiotics in both hospital and non-hospital settings. This bacterial resistance covers almost all antibiotics, including the carbapenems used for treatment against infections caused by these MDR bacteria^1,2^. This problem of antibiotic resistance in human medicine is constantly highlighted by international health organizations such as the CDC (Centers for Diseases Control and Prevention) or the WHO (World Health Organization), which are calling for action to combat this phenomenon^3,4^. This putative antibiotherapy impasse was one of the leading causes of the resurrection in the mid-1990s of abandoned antibiotic, i.e. polymyxin E (colistin), as antibiotic of “last resort” against MDR bacteria^5^. Until 2015, colistin resistance was extremely rare as it was not detected and was exclusively linked to chromosomic mutations in genes involved in Lipid A decoration, i.e. *pmrA/B*, *phoP/Q*, *ccrA/B*, *lpxACD*, or *mgrB* genes that resulted in a modification of bacterRaial membranes by adding sugar (phosphoethanolamine (PEtN) or 4-amino-4-deoxy-L-arabinose (L-ara4N)) to the lipid A moiety^5–8^. The colistin resistance mechanism also involves genes encoding phosphoethanolamine transferase (PET) and/or glycosyltransferase proteins that are essential for membrane phospholipid biosynthesis and appear extremely ubiquitous due to their presence in all areas of life including bacteria, archaea and eukaryotes (plants, arthropods)^6,9,10^. In 2015, a new transferable colistin resistance mechanism has been described, i.e. the mobile colistin resistance *mcr*-1 gene, encoding for a PET^11^ which nowadays, along with new variants of this gene, has been described worldwide in a wide variety of bacterial species and is believed to pose a major public health concern as it can be widely disseminated among pathogenic bacteria by horizontal transfer via transposons and recombinant plasmids^12,13^. In fact, this emerging colistin resistance gene has been already reported in almost all pathogens commonly associated with nosocomial infections and in various sources including food, animal, human and environment (water, soil, etc.) samples^6,13^. Nowadays, nine *mcr* gene variants (from *mcr*-1 to *mcr*-9) have been described, thus amplifying the concern and the interest granted to colistin resistance^14^. Interestingly, most of these *mcr* genes have been discovered into bacteria from animal and/or environmental origin^15–19^. Indeed, *mcr*-1 and *mcr*-2 have been suggested being originated from *Moraxella* species^20^, *mcr*-3 from *Aeromonas veronii*^21^, *mcr-4* from *Shewanella*^22^, *mcr*-8 recently described in *Raoultella ornithinolytica*^15^ and *mcr-9* from *Salmonella enterica* <u>serovar</u> Typhimurium^23^. However, despite the large number of reported publications on these mobilized colistin resistance genes, the sources, origins and roles of these enzymes remain uncertain and not yet described. Here, we conducted a massive bioinformatic analysis of 64’628 downloaded bacterial genomes to investigate the presence and putative source of these *mcr* gene variants in order to understand the presumed roles that these enzymes can play in the biosphere and various ecosystems.

## Results

### Homologous sequences of mobile colistin resistance genes in available database

As a starting point, the first MCR-1.1 protein described (NG_050417) was used as query in a BlastP analysis against the NCBI database to fish out a total of 13’658 protein hits with aa identity ranged from 30% to 100% and alignment ≥ 30% (fig. 1A). Results include all MCR-variants (from MCR-2 to MCR-9) with aa identity ranged from 30.82% with MCR-4.1 to 82.66% with MCR-6.1 (fig. 1B). Interestingly, reference MCR variants exhibit almost the same protein size (1’635 aa in average) but show a significant difference in their GC content ranged from 40.1 % for MCR-4.1 to 56.17% for MCR-7.1 (fig.1B), likely suggesting different origins. As shown in Figure 1A, inferred phylogenetic tree with all obtained sequences reveals a total of 494 bacterial genera (only representative genera are shown on this figure) highlighting a high diversity of PET proteins among bacteria. Moreover, among the 13’658 retrieved hits, 90,89% (n=12’410) exhibit aa identity less than 40% with MCR-1 and only 9.11% (n=1’244) had aa identity > 40% with MCR-1. This finding suggests that these PET enzymes are not specifically hosted by some bacterial genera but are present in a wide range of bacterial species, demonstrating that these latter are extremely ubiquitous in microorganisms.

**Figure 1:**
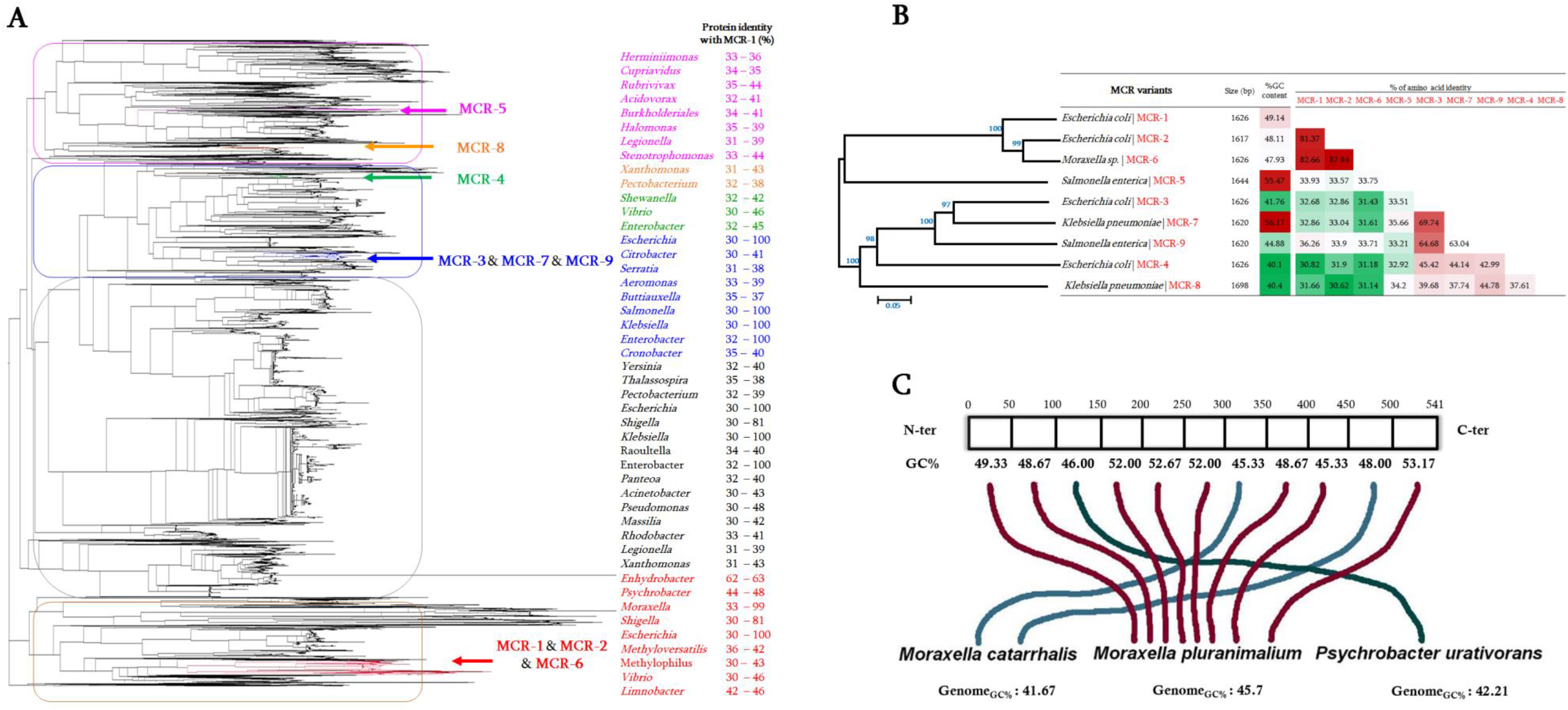
General representation of the distinctive homologous MCR sequences. **(A)** Phylogenetic tree inferred from PETs sequences from different bacterial species and highlighting MCR-1 to MCR-9 group using Archaeopteryx software. Protein identity of MCR-1 with all homologous sequence is indicated, lowest and highest % identity are indicated for each genus. Only representative bacterial genera are shown on this figure. **(B)** General features of known MCR variants including their phylogenetic relationship, nucleotide size and %CG content. Pairwise comparison of amino acid identity of MCR variants is also presented. **(C)** Rhizome analysis of MCR-1. This latter is here split into fragment of 50 aa and phylogenetic tree inferred for each fragment. Each presented line (blue and red) refer to the putative closest ancestor of each corresponding fragment. 8 out of the 11 fragments appeared from *M. pluranimalium* species.

### Rhizome analysis of MCR-1 sequence

To investigate the origin of MCR-1 protein, this latter was split into 11 fragments (50 aa of size) and each fragment was blasted against the NCBI database to retrieve the 100 best BlastP hits. Then, phylogenetic tree analysis was assessed for each fragment to determine its origin. As shown in Figure 1C, 8 out of the 11 MCR-1 fragments appear to be from *Moraxella* genus, especially *Moraxella pluranimalium*.

### Subtree analysis of each MCR variants

As seen on Figure 1A, the phylogenetic tree analysis shows that the 9 MCR variants appear in different branches of the tree. So, subtree containing each MCR variant was retrieved for further detail analysis:

### MCR-1, 2, and 6

These three MCR sequences share more than 80% of identity (fig. 1B) and therefore appear clustered together on the inferred phylogenetic tree (fig. 1A). Therefore, as shown in Figure 2A, the phylogenetic subtree of MCR-1, 2 and 6 reveals that the putative progenitors of these MCR variants would be *Moraxella*, *Enhydrobacter*, *Dichelobacter*, *Psychrobacter*, *Methylophilaceae*, *Limnobacter*, and *Vibrio*. The aa identity with respect to MCR-1 ranged from the lowest 37% with sequences from *Vibrio* to the highest 59% with those from *Moraxella*, *Enhydrobacter* and *Dichelobacter*. It is interesting to note that all these bacteria originate from water, the environment and/or the soil (suppl. Table S1).

**Figure 2:**
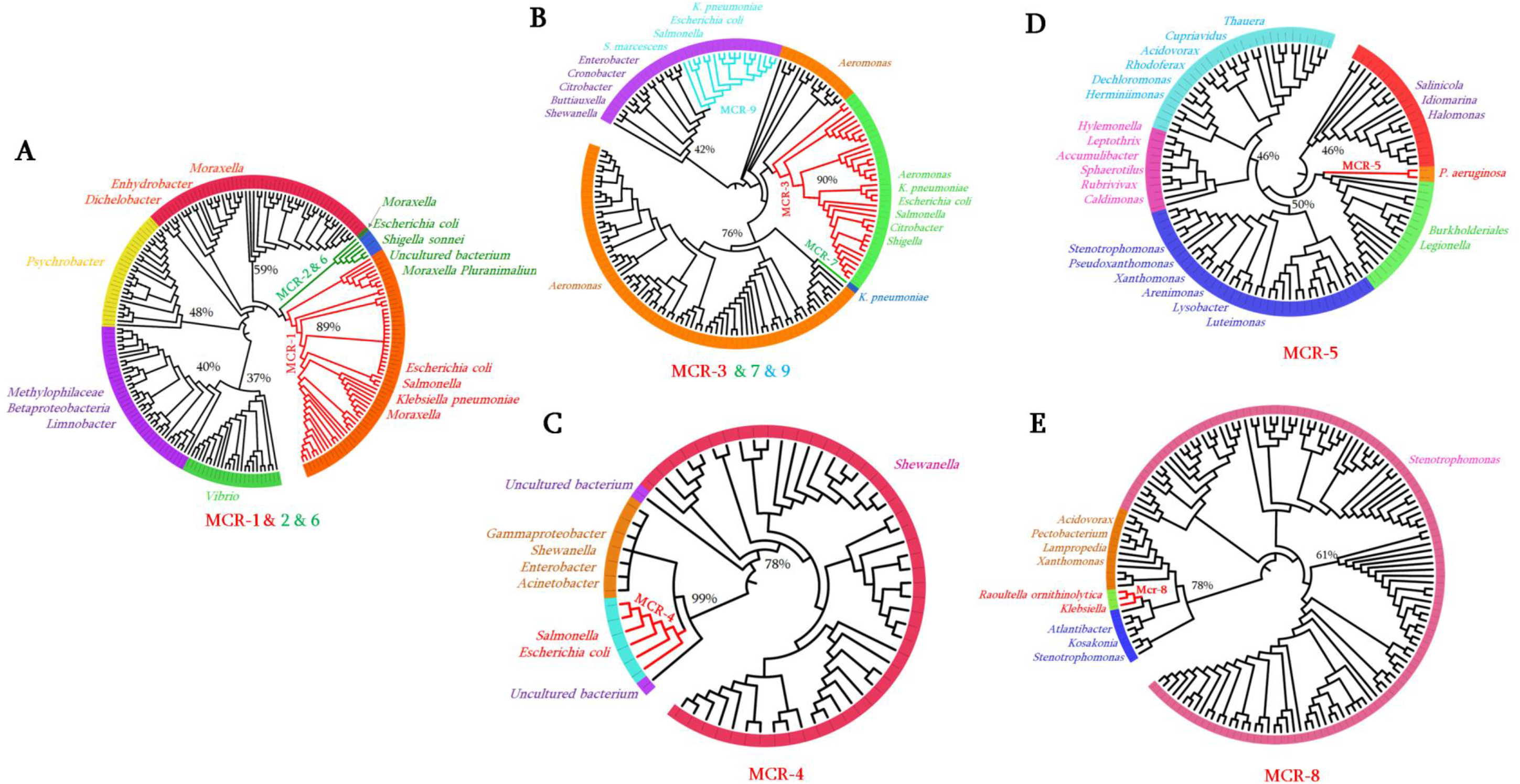
Phylogenetic subtrees highlighting the position of known MCR variants relative to the other species. **(A)** Phylogenetic subtree of MCR-1; MCR-2 and MCR-6; **(B)** Subtree of MCR-3, MCR-7, and MCR-9; **(C)** subtree of MCR-4; **(D)** subtree of MCR-5; and **(E)** subtree of MCR-8. The average percentage of aa identity with MCR variants are indicated.

### MCR-3, 7, and 9

The three sequences share between 63.04 and 69.74% of aa identity despite the significant difference of their %GC content (fig. 1B). The retrieved phylogenetic subtree containing the three variants reveals (fig. 2B) distinct clades of sequences where the most closely related PET sequences were from *Aeromonas* species, well described as water source bacteria. MCR-3 is identified in different *Enterobacteriaceae* species and in *Aeromonas* (fig. 2B). Interestingly, MCR-7 appears on a distinct clade from this tree and shares in average 76% aa identity with *Aeromonas* sequences. MCR-9 sequence appears to be present in *Enterobacteriaceae* species from diverse bacterial genera, including *Serratia, Escherichia, Salmonella*, and *Klebsiella* (fig. 2B). As shown in this figure, MCR-9 shares in average 42% aa identity with sequences from *Cronobacter*, *Enterobacter*, *Shewanella* and *Buttiauxella*; these latter being described with isolation source of water, soil, environment and animals (Suppl. Table S2).

### MCR-4

This MCR variant exhibits the lowest %GC content (40.1 %GC) and shares only 44.14% of aa identity with the most closely related MCR sequence, i.e. MCR-7 (fig. 1B). As shown in Figure 2C, although it has been reported in *Enterobacteriacea*e such as *E. coli* and *Salmonella*, MCR-4 sequence clearly originates from *Shewanella* species with sequences sharing on average 78% of aa identity.

### MCR-5

Homologous sequences to MCR-5 appear extremely diverse and present in various bacterial genera (fig.2D; suppl. Table S1). Interestingly, as *Pseudomonas* genus where MCR-5 has been already reported^24^, all bacteria from this subtree (fig. 2D) have been originating from water, soil and environment (Suppl. Table S1). The most closely related sequences to MCR-5 from this tree were identified in *Legionella* and *Burkholderiales*.

### MCR-8

MCR-8 is highly distant to the other variants. This latter has so far only been reported in *K. pneumoniae* and *R. ornithinolytica*; and appears to be closely related to PET sequences from *Atlantibacter*, *Kosakonia*, *Xanthomonas*, *Lampropedia*, *Pectobacterium*, *Acidovorax*, and *Stenotrophomonas* (fig. 2E). *Stenotrophomonas* exhibits a large number of homologous MCR-8 sequences, thus suggesting as the origin of this MCR variant.

Thus, based on these results, which show different bacterial origins and hosts of these MCR sequences, we have investigated the nine MCR variants in all available sequenced bacterial genomes with clinical interest and those from water and environment sources in order to identify the putative reservoirs of these enzymes.

### MCR variants in sequenced bacterial genomes

A total of 64’628 Gram-negative bacterial genomes were downloaded from the NCBI RefSeq database and include 60 bacterial genera constituted by 1’047 bacterial species (fig. 3). As presented in this figure, a RpoB-based phylogenetic tree was inferred to establish their phylogenetic relationship. Only 4’214 out of the 64’628 genomes (6.52%) were complete genomes and the remaining genomes were a Whole-genome sequences (WGS). BlastP analysis of the nine MCR sequence variants against all downloaded data set provided significant positive results for MCR-1.1, MCR-2.1, MCR-3.1, MCR-4.1, MCR-5.1, MCR-8.1 and MCR-9.1 with aa identity values ranging from 91 to 100% and alignment ≥98%. A total of 1’386 MCR hits were identified in 15 out of the 60 genera analysed. As shown in Figure 3, 952 MCR-1 hits were identified, including 862 hits in *E. coli* genomes, 43 hits in *K. pneumoniae*, 31 hits in *Salmonella* spp., 15 hits in *Shigella* spp. and one hit in *Acinetobacter baumannii*. MCR-2 was detected in only two genomes (*M. pluranimalium* and *S. sonnei*) (fig. 3). MCR-3 was detected in diverse bacterial genera and was interestingly detected in all the *Aeromonas* species analyzed and almost in all *Shigella* species. As found in our first analysis (fig. 2B), *Aeromonas* species appears clearly as the bacterial reservoir of MCR-3 variant (fig. 3). Six MCR-4 hits were detected in our analysis, including 1 hit *in Enterobacter cloacae*, 2 hits in *Shewanella* spp. and 3 hits in *A. baumannii* and *A. nosocomialis*. MCR-5 was identified in only five species, including *E. coli* (n= 11 hits), *S. enterica* (n=3 hits), *A. salmonicida* (n= 1 hit), *Cupriavidus spp*. (n= 1 hit) and *P. aeruginosa* (n= 1 hit). MCR-8 was identified in only *K. pneumoniae* (n= 20 hits) and its related species i.e. *R. ornithinolytica* (n= 2 hits), previously classified in *Klebsiella* genus. It is interesting to note that the newly described MCR-9 from *S. enterica* serotype *Typhimurium* strain isolated from a patient in Washington in 2010, was the most distributed sequence among bacterial species and was identified in 309 genomes (aa identity ≥ 99.81% and 100% of alignment) representing 10 out of the 14 MCR-positive genera (fig. 3). Common *Enterobacteriaceae* species appear as the reservoir of this gene, especially *E. cloacae*, *S. enterica*, *K. pneumoniae*, and *E. coli*, where 134, 72, 44, and 24 hits were identified respectively. Moreover, we observed for the first time the presence of MCR-9 variant in *Serratia marcescens* and *Proteus mirabilis* genomes, those being naturally resistant to colistin.

**Figure 3:**
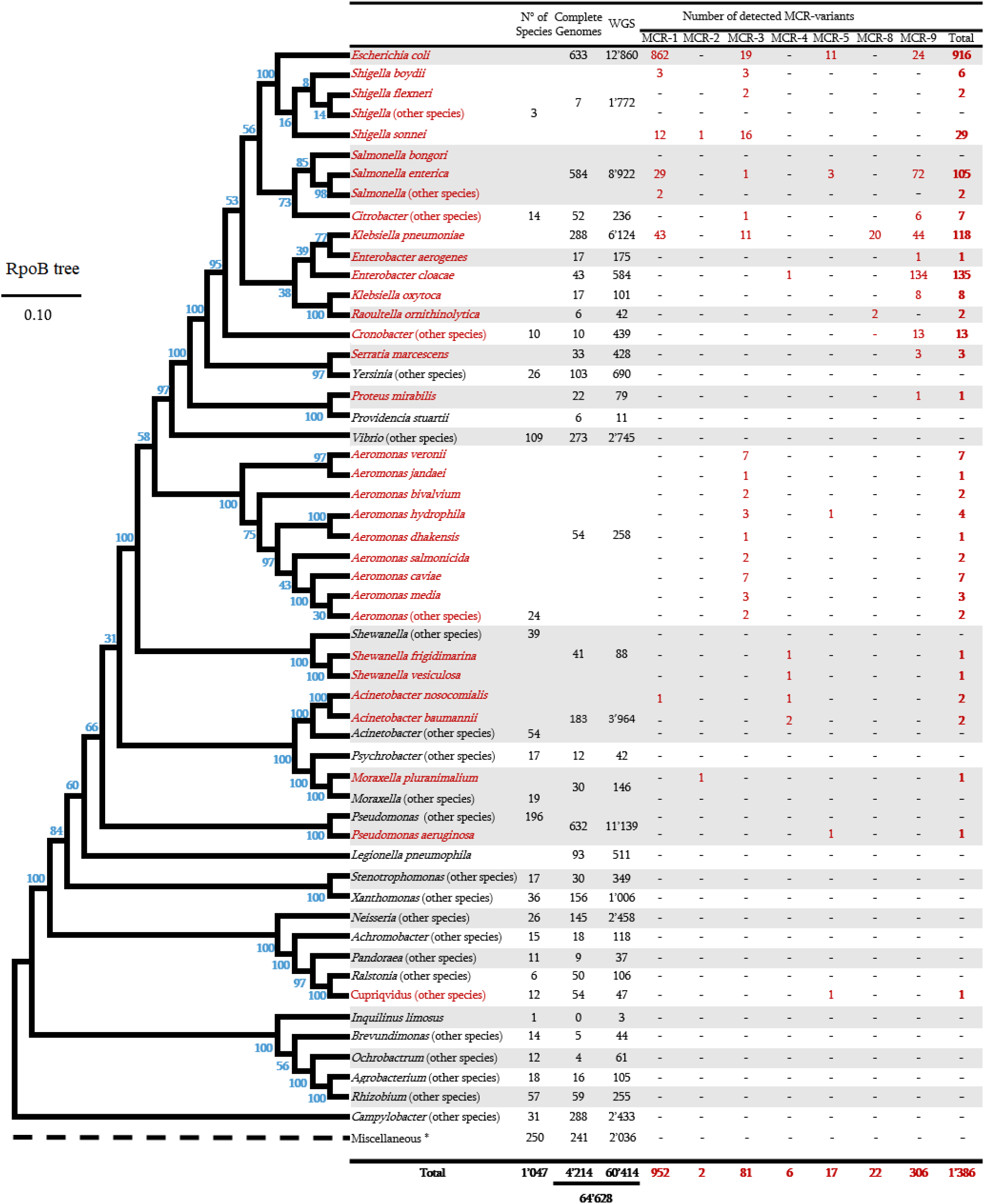
All bacterial genomes analysed here and the RpoB-based phylogenetic tree of all investigated bacterial species. Bacterial species identified as MCR-producers and the number of significant MCR hits are highlighted in red. The number of species and genomes analyzed (complete and Whole genome sequences) are also indicated. *, refers to rare bacterial species from the environment (list of the 27 genomes are given in **Suppl. Table S1**).

Except MCR-1-producing *E. coli* strains that have been well described^25,26^, detailed metadata of all MCR-producer strains including strain names, aa identity with MCR variants, Bioproject number, collection dates, geographical location, isolation sources, and host organisms are given in Suppl. Table S2.

Apart from these significant occurrences (aa identity ≥ 90% and alignment ≥98%) with reference MCR variants (from MCR-1 to MCR-9), we searched for new MCR candidate sequences using threshold values of 50% ≤ aa identity ≥90% and alignment ≥90%. According to that, 5’265 additional hits were identified and distributed among 25 genera including among others *Escherichia, Aeromonas, Vibrio, Stenotrophomonas, Moraxella, Klebsiella, Salmonella, Shewanella, Shigella, Enterobacter, Raoultella, Serratia, Citrobacter, Xanthomonas, Acinetobacter* and *Proteus* (Suppl. fig. S1). Interestingly, 1’244 out of the 5’265 hits exhibit aa identity between 75% and 90% with MCR variants, those being putative new MCR variants (Suppl. fig. S1B). The *E. coli* species appears in this analysis as the main reservoir of these genes with a total of 3’090 MCR sequences identified (Suppl. fig. S1). Surprisingly, a huge number of MCR-9-like sequences were identified with a total of 3’056 hits. Moreover, most of these bacterial genera including, *Stenotrophomonas*, *Vibrio*, *Aeromonas*, *Shewanella*, *Moraxella*, *Buttiauxella*, and *Salmonella* are bacteria of environmental sources.

As shown in Suppl. Figure S2, all identified MCR hits from analysed genomes i.e. the 1’386 significant hits (aa identity ≥ 90%) and the 5’265 additional hits (50% ≤ aa identity ≥ 90%) were pooled together to infer a phylogenetic tree. We noticed that new putative MCR sequences (in blue) significantly clustered with the significant described MCR variants (in red). This suggests the presence of real MCR candidates that may soon emerge in common pathogenic bacteria. Moreover, our analyses reveal that all the analysed 64’628 genomes have constitutive PET sequences that exhibited an identity between 30% and 50% with MCR variants (data not shown).

### Genomic environment of *mcr* variant genes in bacterial genomes

As previously described, mobilized colistin resistance genes are known to be transmitted between bacteria through mobile genetic elements (transposons and plasmids). Except *mcr*-1 gene that has been well described in the literature regarding to his transmission mechanism^11^, we investigated, from downloaded genomes, the genetic environment of the other *mcr* genes to figure out their mode of transmissions.

As shown in Figure 4A, while *mcr*-2 exhibits a different genetic structure in the two genomes identified, *mcr*-3 gene appears in a same sequence composed by it-self and a *dgkA* gene encoding for diacylglycerol kinase protein in four different bacterial species (fig. 4B) and was associated with an insertion sequence (IS) element upstream or/and downstream to *mcr*-3. Genome analysis reveals that this genetic structure of *mcr*-3 in *K. pneumoniae* was identical in the 11 genomes positive for *mcr*-3. For *Aeromonas*, the *mcr*-3-containing transposon was also identical in 26 out of the 28 genomes positive for *mcr*-3. The 21 genomes of *Shigella* also exhibited the same *mcr*-3-containing transposon. Interestingly, in the 19 *E. coli* genomes, only two of them exhibited the structure presented in this Figure 4B.

**Figure 4:**
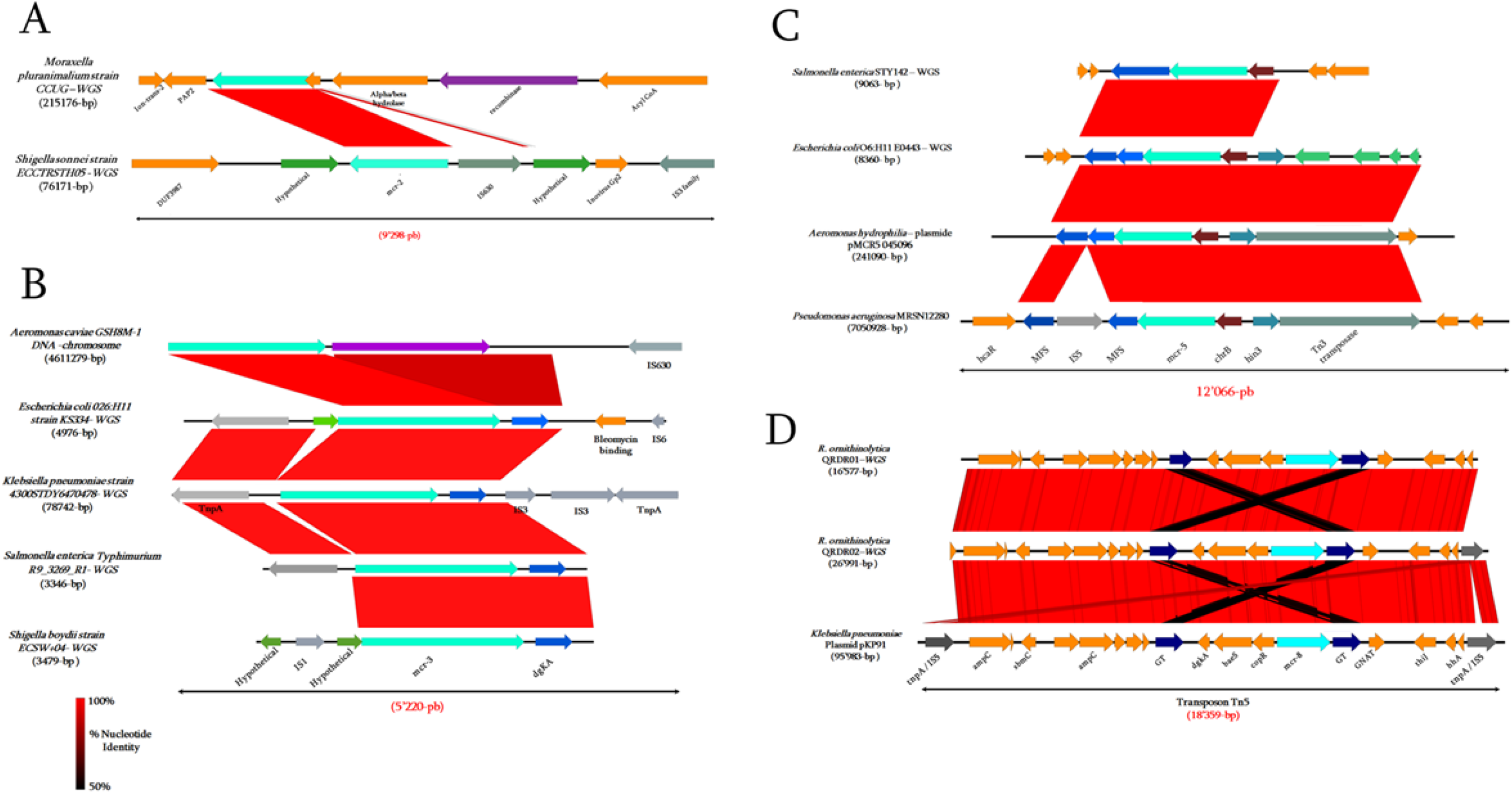
Genetic environment analysis of *mcr-2*; *mcr-3*; *mcr-5* and *mcr-8* in bacterial genomes. **(A)** Comparison of the *mcr-2* identified in *M. pluranimalium* and *S. sonnei*. **(B)** Genetic environment of *mcr-5* identified in. *S. enterica*; *E. coli*; *A. hydrophilia* and *P. aeruginosa*. **(C)** Genetic environment of *mcr-3* identified in *A. caviae. E. coli. K. pneumoniae. S. enterica Typhimurium* and *S. boydii*. **(D)** Comparison of the *mcr*-8 identified in *K. pneumoniae* and *R. ornithinolytica*. The arrows indicate the positions and directions of the ORFs. Regions of more than 90% homology are marked by red shading.

The *mcr*-5-containing transposon presented in Figure 4C appears in the same structure in the 11 *E. coli* genomes and the 3 *Salmonella* genomes. We also observed that this *mcr*-5 gene is associated in this mobile genetic element with transposase and IS elements IS5/Tn3 in the different genomes. This genetic organization suggests sequence acquisition by sequence recombination via these transposases and IS elements in these genomes.

Among the 20 positive genomes of *K. pneumoniae* and 2 of *R. ornithinolytica* for *mcr*-8 gene, the Tn5 transposon (fig. 4D) had the same structure in 13 out of the 22 genomes. This transposon located on conjugative plasmid in both species was highly similar and bordered in both sides by transposase *tnpA*/IS903 (fig. 4D). Interestingly, *mcr*-8 gene appeared associated with *copR*, *baeS* and *dgkA* genes encoding respectively for a transcriptional activator protein CopR, an integral membrane sensor signal transduction histidine kinase and a diacylglycerol kinase. Moreover, we identified two glycosyl transferases (GT) upstream and downstream of the *mcr*-8 gene and two *ampC* serine-β-lactamase genes. This genetic organization also suggests sequence acquisition of the transposon Tn903 by horizontal transfer from exogenous origin.

Finally, the transposon containing *mcr*-9 shown in Figure 5 has the same structure in the nine genomes analysed, except in *P. mirabilis*. This transposon was highly similar and bordered in both sides by a transposase *IS5*/IS6. We identified a *wbuC* family gene encoding for cupin fold metallo-protein, which was highly conserved in this transposon. Moreover, a conserved nickel/cobalt operon (i.e. *rcnA, rcnR, pcoE* and *cusS* genes), which plays an important role in copper tolerance under anaerobic growth and nickel homeostasis in bacteria was identified in almost all analysed transposons (fig. 5). Some specific sequences including transcription factors (e.g. Ni(II)/Co(II)-binding transcriptional repressor RcnR, response regulator, transcriptional regulator ArdK, and Sensor kinase CusS), antimicrobial resistance genes (phosphotransferases *(aph(3”)-Ib, aph(6)-Id)*, copper-binding protein PcoE and zinc metalloprotease Zinc M) were identified in these transposons (fig. 5). This genetic organization suggests also sequence acquisition by transposition. Interestingly, *mcr*-9 gene is strongly associated with *wbuC* gene encoding for Cupin fold metallo-protein located upstream in the different transposon of different bacterial species. This may suggest an essential role of this *wbuC* gene for the activity of MCR-9 enzyme.

**Figure 5:**
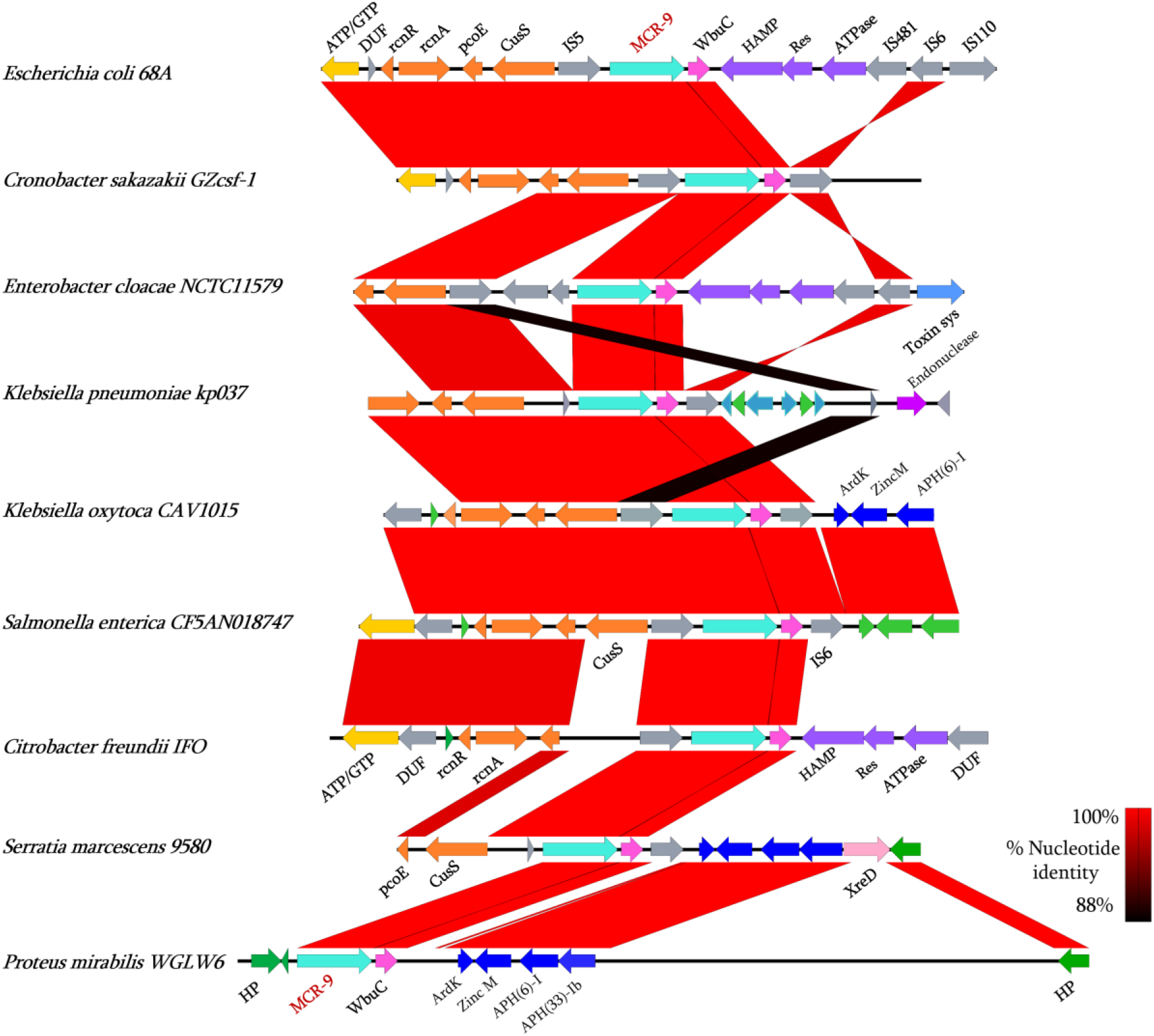
Genetic environment of *mcr*-9 gene in different bacterial genomes of different bacterial species. Gene size, % GC content and predicted function are indicated in **suppl. Table S3.**

### MCR-4 variant in *Acinetobacter* species

So far, MCR-4 has only been described for this bacterial species in a genome of *A. baumannii*. Here, we identified three genomes of *Acinetobacter* (two *A. baumannii* and one *A. nosocomialis*) containing *mcr*-4 gene variant. The two *A. baumannii* strains (AB18PR0652 and MRSN15313) were isolated respectively from feces from China in May 2018 and from a cerebrospinal fluid from Brazil (Salvado) dating from 2008, while the latter genome has not been described or analyzed in the literature (Suppl. Table S2). In both isolates, the *mcr*-4 gene was located on plasmid pAB18PR065 and pAB-MCR4.3, respectively. As shown in Figure 6A, these two plasmids were compared to that identified in *A. nosocomialis* isolate T228 and appear very similar. Moreover, the six *mcr*-4-containing transposons identified here were compared (fig. 6B). As shown in this figure, these transposons were similar only between *Acinetobacter* species and were integrated into these plasmids by splitting an HNH endonuclease encoding gene (fig. 6B). In addition, a recombinase and two toxin/antitoxin systems were identified in *Acinetobacter* and *Shewanella* on the *mcr*-4-harboring-transposon.

**Figure 6:**
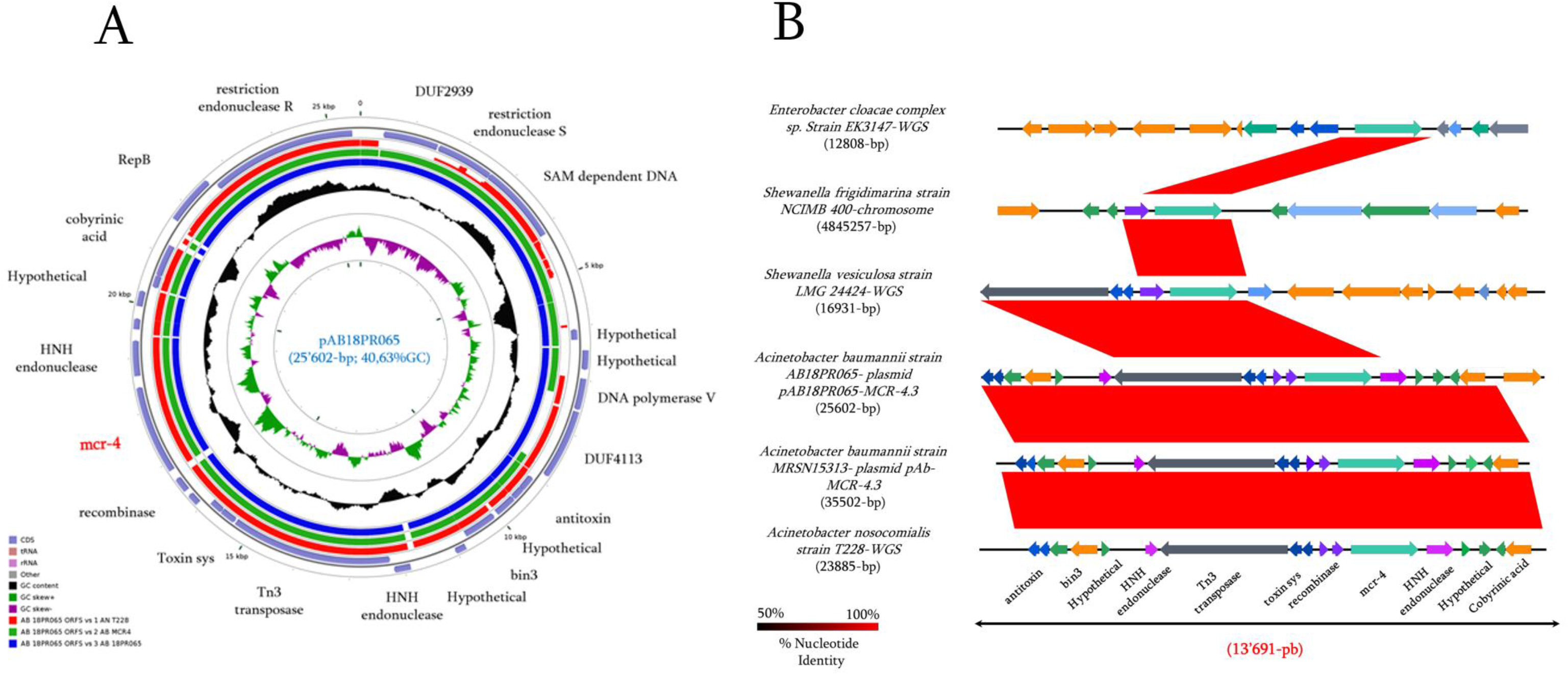
Comparison of the genetic environment of *mcr*-4 gene in bacterial genomes. **(A)** Comparison of the three plasmids harboring *mcr*-4 gene identified in *A. baumannii* and *A. nosocomialis* using CGView software. **(B)** Genetic environment of the *mcr*-4 gene. Gene size, % GC content and predicted function are indicated in **suppl. Table S3.**

### Metadata analysis of all MCR positive isolates in analysed genomes

The presented metadata in the suppl. Table S2 were subjected to cytoscape analysis in order to highlight links between isolates, MCR variants and isolation sources. So, as shown in Figure 7, according to the isolation sources, almost all MCR variants were detected in bacterial species isolated from the three investigated sources (i.e. Human, Animal, and Environment).Three bacterial species including *K. pneumoniae*, *E. cloacae*, and *S. enterica*, all isolated from Human, appeared as the most predominant bacterial species harboring MCR enzymes. MCR-3 was most detected variant in different species and was also mostly detected in isolates from Animals. Interestingly, only MCR-9 variant was detected in bacterial strains isolated exclusively from human and these isolates were almost all *Enterobacteriaceae* species (fig. 7). These findings suggest that MCR-9 variant can became soon the most concern in human medicine regarding the colistin resistance.

**Figure 7:**
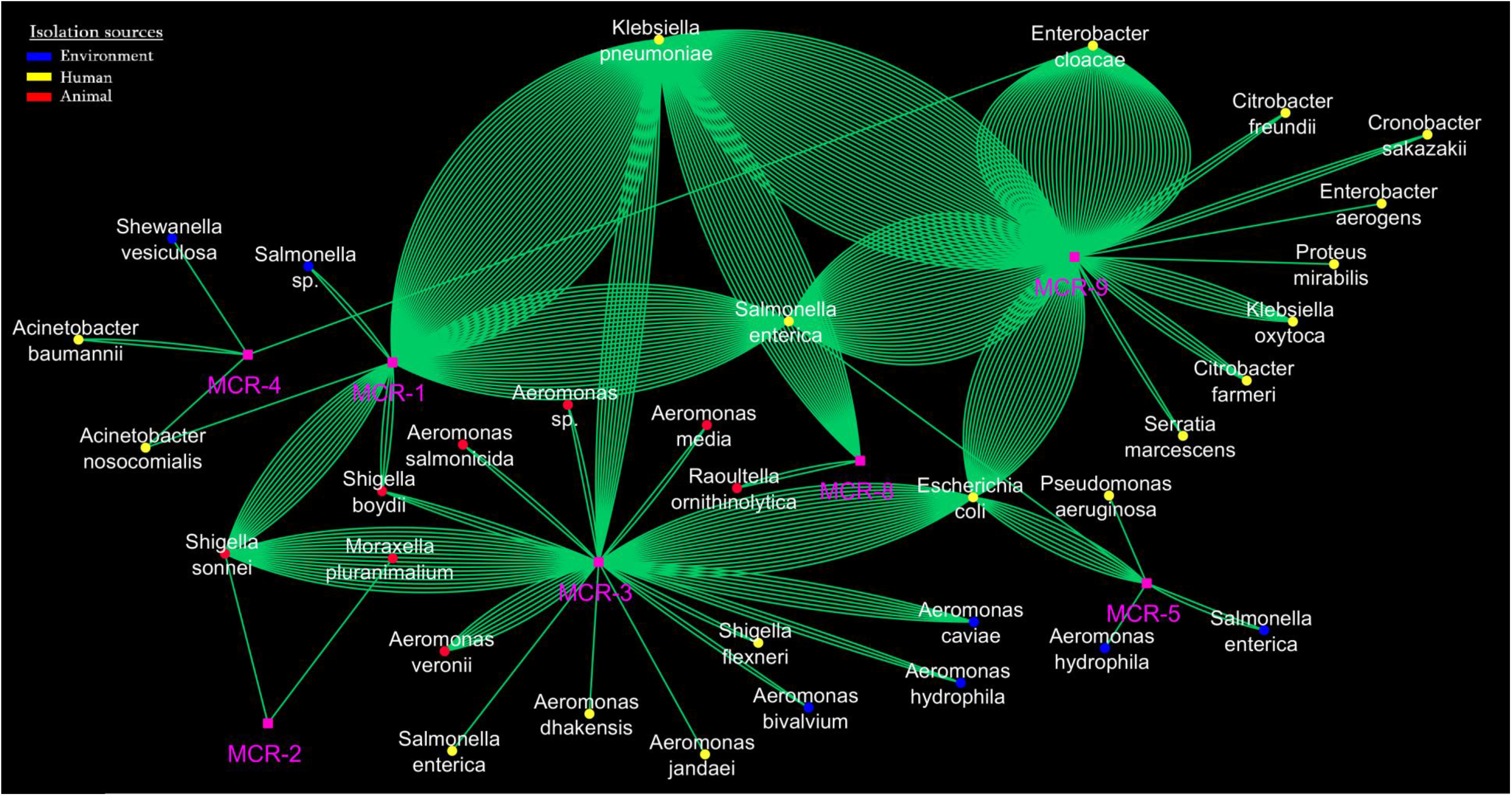
Network analysis of metadata of MCR variant producer isolates (excluding *E. coli mcr*-1) highlighting the link between each MCR variant and the different bacterial species and their isolation sources such as Human, Animal, and the Environment.

## Discussion

Colistin resistance has become an increasing concern for human medicine since the emergence and spread of mobile colistin resistance genes^11,13^. The presence of these genes worldwide in all ecosystems including animals, environment (soil, water, plant) and humans^27^ raises the question of their sources, origins and roles in a global context of reservoir of antibiotic resistance genes. We show here how a large-scale data analysis approach, by studying more than 60’000 genomes, can help us to understand the bacterial reservoir and source of a gene family. We figured out that almost all mobile colistin resistance genes (from MCR-1 to MCR-9) described originated from environmental bacteria, especially from water sources. Indeed, as proof of concept, the rhizome analysis performed reveals the origin of MCR-1 from *M. pluranimalium*, as previously suggested by another approach^28^. Moreover, our findings reveal that PET genes are extremely ubiquitous in bacteria since detected in 1’047 species analyzed here. Our massive genome analysis approach allowed us to identify the source and to detect the presence of these MCR-like sequences from available bacterial genomes. This was the case for MCR-3 and MCR-7 originating from *Aeromonas* spp. Interestingly, among these MCR-3-like sequences, one of them identified from *Salmonella* and *Buttiauxella* spp. has recently been published as the new MCR-9 variant^29–32^. We identified *Shewanella* spp. as progenitor of MCR-4 sequences, that have been detected in *Acinetobacter* spp., also recently reported^33^. MCR-8 newly reported in *Raoultella* spp.^15^ was also identified and appears to be present in *Stenotrophomonas* spp. As a confirmation of this hypothesis, this latter has been recently described in *Stenotrophomonas* strain from sewage water^34^. Interestingly, whilst MCR-8 has only been reported in less than five studies to date, we identified 22 MCR-8 in already sequenced genomes of *Klebsiella* and *Raoultella* spp. All these MCR variants newly described or in progress to be published are real true witnesses of our results which predict putative 5’265 MCR-like sequences that could emerged soon in common pathogenic bacteria.

Interestingly, we observed in our study that most of the bacteria analysed are from water sources as highlighted in the Suppl. Table S1. This suggests that water environment appears to be the main reservoir and source of these MCR-like genes. Indeed, almost all MCR variants described so far have been reported from bacteria from water in different environments. MCR-1 was detected and identified directly from swine wastewater of purification facilities^35,36^, directly from water or from bacteria (e.g. *E. coli* and *E. cloacae*) collected from urban surface water, rivers or lakes^37–39^. This gene was reported from wastewater treatment plants in China^40^, in South Africa^41^, in Italia^42^, in Germany^43^ or from hospital sewage water in China^44^. This variant was also reported from various bacteria isolated from other water environments, such as seawater in Algeria^45^, from top water in rural China^46^ or from freshwater of intensive aquaculture in China^17^. MCR-3 was reported from *Aeromonas* species isolated from river water^37^. It has recently been reported that the aquatic bacteria of the genus *Shewanella* are a reservoir/progenitor for *mcr*-4 encoding gene^47^. MCR-5 and MCR-8 were described from the same *Stenotrophomonas* strain isolated from sewage water in China^34^. Overall, urban wastewater treatment plants are one of the most common reservoir and source of antibiotic resistance genes (ARGs) against various antibiotic classes (β-lactams, aminoglycosides, quinolones, tetracyclines, sulfonamides, macrolides)^48^. Of course, these AR genes, including MCR-like genes, were also reported from other sources including plants, soils^49^ and various animal species^49,50^, most of them being in direct contact with water. Therefore, according to this, aquatic environment represents a great source and reservoir of MCR-carrier bacteria. Obviously, the global overuse of antibiotics in human medicine, animals and agriculture, has a great impact on the selection and the emergence of ARGs and multidrug resistant bacteria but this could not explain that more than 10% of bacterial genomes analyzed in our study contain a PET gene.

Indeed, our results revealed that PET genes, including *mcr-*like genes, are extremely ubiquitous in bacteria and in all areas of life^6,9,10^, thus raising the question of their real roles. It is unlikely that colistin pressure in the environment and animals, especially in water environment, may be the only cause of the existence of such a ubiquitous presence of these enzymes that could be mobilized within bacteria. This raises the question of the other roles and functions these enzymes may have in nature and in the biosphere. Because polymyxins share chemical similarities with cationic antimicrobial peptides (cAMPs) secreted by vertebrates and invertebrates, it is possible that decoration of lipid A of those bacteria with ethanolamine constitutes an immune defense against those metabolites. There is evidence that expression of PET in *Haemophilus ducreyi* is associated with polymyxin resistance as well as resistance to human defensins^51^. Although this hypothesis may explain another role of these enzymes against innate immunity from eukaryotic cells, this could not explain the huge prevalence of those enzymes in water environments. Interestingly, the most common cell living in water sources are bacteriophages^52,53^. Predation of bacterial cells by bacteriophages is linked to the interaction between phages and O-antigens structures of Gram-negative bacteria^54^. One may speculate that decoration of lipid A with ethanolamine is a “camouflage” of the antigen receptor of bacteria against bacteriophages. This defense hypothesis against bacteriophage can be supported by a recent study reporting a disruption of *mcr*-1 gene by a bacteriophage P1-like in an atypical enteropathogenic *E. coli* recovered from China^55^. Thus, we believe that PET genes in bacteria, by adding sugars to the bacterial cell membrane constitutes a very ubiquitous and old immune defense system that bacteria have developed over time, not only to resist to polymyxins, but also to resist to eukaryotic innate immune system and bacteriophage predation.

Finally, our innovative approach, i.e. massive genome analysis, appears to be a powerful tool and a new concept that opens a new field of big data analysis and research; *mcr* gene variants are here only an example but this approach can be carried out to explore the source and the origin of any gene family. PET genes such as *mcr* genes are extremely ubiquitous in bacteria mainly from the environment, soil and especially from water environments. These latter could be mobilized by transposable elements and transmitted by horizontal transfer to bacteria as reported in the literature since the emergence of *mcr*-1 in 2015. We believe that those enzymes play other major roles as immune defense systems in the biosphere that should be further explored in the future because this may have a great impact, not only for human health, but also as a one health perspective.

## Material and Methods

### Looking for homologous MCR-1 sequences from the NCBI database

Reference sequence of MCR-1 (AKF16168.1) was used as query for BlastP analysis against the NCBI database using as threshold e-value 10e-5. All homologous sequence with identity ≥30% and alignment ≥30% were kept for further analysis. Protein alignments were performed using Mafft tool (https://mafft.cbrc.jp/alignment/software/) and phylogenetic trees were inferred using neighbor-joining method in FastTree program (http://www.microbesonline.org/fasttree/) and visualized with FigTree v1.4.2 (http://tree.bio.ed.ac.uk/software/figtree/) and Archaeopteryx 0.9920 (http://en.bio-soft.net/tree/ATV.html). A rhizome analysis^56^ was carried out for MCR-1 to investigate putative ancestral recombination events of bacterial sequences leading to creation of chimerical gene. For this purpose, protein sequence of MCR-1 (AKF16168.1) was split into 10 fragments of 50 amino acids. Each fragment was blasted against the NCBI database using as threshold e-value 10e-5 to select the 100 best homologous sequences according to the aa identity ≥ 50% and coverage ≥80%. From these sequences, phylogenetic tree with each fragment was constructed to infer the closest ancestor.

### Looking for all MCR variants in bacterial genomes

Whole genome sequences of the most common Gram-negative bacteria were retrieved from RefSeq data: NCBI Reference Sequence Database (ftp://ftp.ncbi.nlm.nih.gov/genomes). This access to the repository is made through version 0.2.7 of the NCBI genome download scripts, which allowed the download of the entire genome assembly, including all required data for each genome, genomic sequences, annotated proteins and assembly structure reports. Reference protein sequence of each MCR variant was also retrieved from the NCBI database. To the best of our knowledges, today (July 10, 2019), nine MCR variants have been described including MCR-1, MCR −2, MCR -3, MCR -4, MCR -5, MCR -6, MCR -7, MCR-8 and MCR -9 with respectively 18, 2, 29, 6, 3, 1, 1,2 and 1 subvariants. BlastP analysis was conducted to investigate the presence of the nine MCR variants (i.e. MCR-1.1, MCR-2.1, MCR-3.1, MCR-4.1, MCR-5.1, MCR6.1, MCR-7.1, MCR-8.1 and MCR-9.1) in the large downloaded data. MCR hits were considered significant on the basis of 90% aa identity and 98% of alignment. Reference sequences of MCR variants were obtained from the Pathogen Detection database (https://www.ncbi.nlm.nih.gov/pathogens/).

### Analysis of genetic environment of *mcr* gene variants

Genetic environments of *mcr* genes were investigated in the selected genomes using the Easyfig version x2.2.3 program (http://mjsull.github.io/Easyfig/) and CGView program (http://wishart.biology.ualberta.ca/cgview/). An annotation using Prokka (http://www.vicbioinformatics.com/software.prokka.shtml) and BlastP were made for genetic environment verification.

### Metadata and network analysis

Metadata of each bacterial genome containing significant hits of *mcr* gene variants (≥ 90% identity, ≥ 98% alignment) including bacterial strain name, bioproject number, collection date, geographical location, isolation source and host organisms, were collected from PATRIC database (https://www.patricbrc.org/) and detailed in **Suppl. Table S2**. To investigate putative links between bacterial strain harboring *mcr* gene variants, metadata information are subjected to networking analysis using the cytoscape software v.3.7.

## Acknowledgements

Financial support from the IHU Mediterranee-Infection, Marseille, France; CookieTrad for English corrections of the manuscript are gratefully acknowledged.

## Author contributions

J.-M.R. and S.M.D conceived and designed the study. M.B.K, S.M.D., S.A.B, T.R., R.R. and J.-M.R. analysed and interpreted data. M.B.K, S.M.D, J.-M.R. drafted the manuscript. S.M.D., S.A.B, R.R. and J.-M.R made corrections and critical revisions of the manuscript. All of the authors read and approved the final version of the manuscript.

## Ethical issue

Not required

---

**Suppl. Figure S1:**
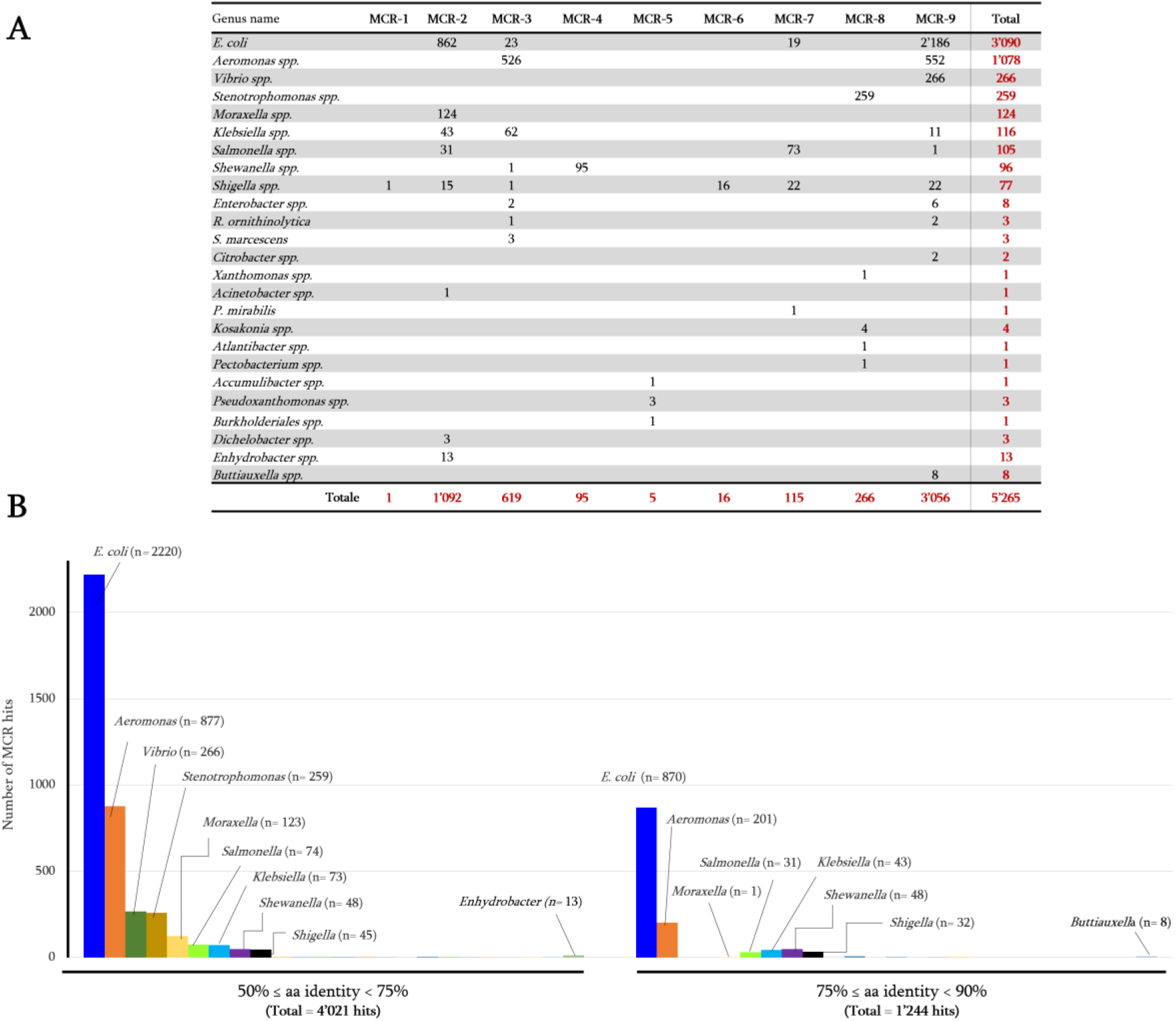
Distribution of MCR hits with aa identity between 50% to 90% with all MCR reference variants. (**A**) Table showing the distribution of the total 5’230 hits among the 16 out of the 32 bacterial genera analyzed. (**B**) Graph showing the distribution of these sequences according to bacterial genera and to aa identity between 50 and 75% and those with identity from 75 to 90%.

**Suppl. Figure S2.**
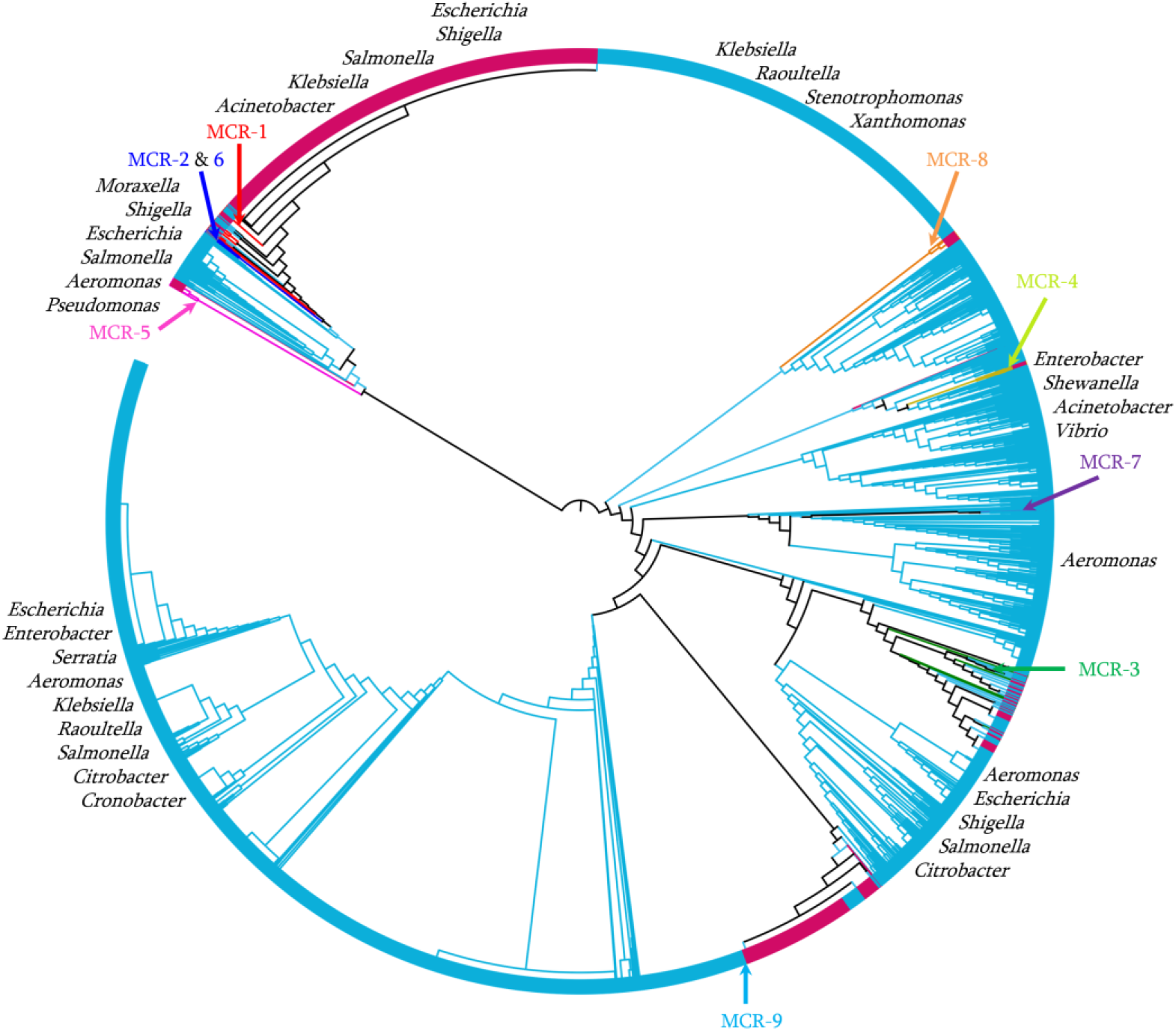
Phylogenetic tree performed from MCR variant hits (n=6’648 proteins) with aa identity between 50% and 100% and alignment ≥90%.

