## Supplementary Table S1 for "Massive analysis of 64’628 bacterial genomes to decipher a water reservoir and origin of mobile colistin resistance (*mcr*) gene variants: is there another role for this family of enzymes?"

**Suppl. Table S2:** MCR variants hits of available bacterial genomes from the different subtrees presented in Figure 2.

| Genus name | Source | MCR variants | Species | Complete Genomes | WGS | GC% | MCR-1 | MCR-2 | MCR-3 | MCR-4 | MCR-5 | MCR-6 | MCR-7 | MCR-8 | MCR-9 | Total |
|---|---|---|---|---|---|---|---|---|---|---|---|---|---|---|---|---|
| ***Enhydrobacter*** | Sea water | MCR-1, 2 & 6 | 2 | - | 15 | 43.6 | - | 12 | - | - | - | - | - | - | - | **12** |
| ***Dichelobacter*** | Sea water | MCR-1, 2 & 6 | 1 | 1 | 2 | 44.4 | - | 3 | - | - | - | - | - | - | - | **3** |
| ***Methylophilaceae*** | Freshwater | MCR-1, 2 & 6 | 24 | 6 | 62 | 50.3 | - | 57 | - | - | - | - | - | - | - | **57** |
| ***Limnobacter*** | Sea water, environment, soil | MCR-1, 2 & 6 | 2 | - | 13 | 52.2 | - | 12 | - | - | - | - | - | - | - | **12** |
| ***Buttiauxella*** | Soil, animal, human | MCR-3 & 7 & 9 | 8 | 1 | 9 | 52.6 | - | - | 1 | - | - | - | - | - | 13 | **14** |
| ***Salinicola*** | Sea water | MCR-5 | 7 | 1 | 17 | 63.6 | - | - | - | - | 3 | - | - | - | - | **3** |
| ***Idiomarina*** | Sea water | MCR-5 | 28 | 5 | 45 | 47 | - | - | - | - | - | - | - | - | 2 | **2** |
| ***Halomonas*** | Sea water | MCR-5 | 59 | 18 | 121 | 55.9 | - | - | - | - | 46 | - | - | - | - | **46** |
| ***Burkholderiales*** | Soil, water, human | MCR-5 | 2 | 124 | 1'383 | 66.4 | - | - | - | - | 2 | - | - | - | 1 | **3** |
| ***Luteimonas*** | Sea water, environment | MCR-5 | 4 | 4 | 7 | 69.3 | - | - | - | - | 5 | - | - | - | - | **5** |
| ***Lysobacter*** | Soil | MCR-5 | 15 | 10 | 34 | 68.3 | - | - | - | - | 34 | - | - | - | - | **34** |
| ***Arenimonas*** | Sea water, environment, soil | MCR-5 | 7 | - | 8 | 70 | - | - | - | - | 5 | - | - | - | - | **5** |
| ***Pseudoxanthomonas*** | Soil, plant | MCR-5 | 7 | 3 | 33 | 69 | - | - | - | - | 42 | - | - | - | - | **42** |
| ***Caldimonas*** | Hot spring (water) | MCR-5 | 2 | - | 2 | 66 | - | - | - | - | 2 | - | - | - | - | **2** |
| ***Rubrivivax*** | Hot spring (water) | MCR-5 | 3 | 1 | 13 | 68.4 | - | - | - | - | 11 | - | - | - | 1 | **12** |
| ***Sphaerotilus*** | Rivers, sewage | MCR-5 | 2 | - | 3 | 69.9 | - | - | - | - | 2 | - | - | - | - | **2** |
| ***Accumulibacter*** | Water | MCR-5 | 3 | 1 | 24 | 62.1 | - | - | - | - | 2 | - | - | - | - | **2** |
| ***Leptothrix*** | Groundwater | MCR-5 | 3 | 1 | 2 | 68.9 | - | - | - | - | 2 | - | - | - | - | **2** |
| ***Hylemonella*** | Wastewater | MCR-5 | 2 | - | 6 | 55.2 | - | - | - | - | 2 | - | - | - | - | **2** |
| ***Herminiimonas*** | Bottled mineral water | MCR-5 | 4 | 2 | 3 | 56.4 | - | - | - | - | 4 | - | - | - | - | **4** |
| ***Dechloromonas*** | Environment, human gut | MCR-5 | 4 | 2 | 9 | 61 | - | - | - | - | 2 | - | - | - | - | **2** |
| ***Rhodoferax*** | Seawater | MCR-5 | 7 | 6 | 6 | 61.4 | - | - | - | - | 4 | - | - | - | - | **4** |
| ***Acidovorax*** | Soil | MCR-5 | 18 | 14 | 73 | 64.8 | - | - | - | - | 25 | - | - | - | 7 | **32** |
| ***Thauera*** | Hot spring (water) | MCR-5 | 13 | 6 | 14 | 66.4 | - | - | - | - | 12 | - | - | - | - | **12** |
| ***Pectobacterium*** | Soil, plant | MCR-8 | 12 | 26 | 110 | 51.8 | - | - | - | - | - | - | - | 48 | 88 | **136** |
| ***Atlantibacter*** | Human, soil | MCR-8 | 2 | 1 | 5 | 54.1 | - | - | - | - | - | - | - | 3 | 1 | **4** |
| ***Kosakonia*** | Environment, soil | MCR-8 | 9 | 8 | 17 | 53.9 | - | - | - | - | - | - | - | 30 | 1 | **31** |
| **Total** | | | **250** | **241** | **2'036** | **-** | **0** | **84** | **1** | **0** | **205** | **0** | **0** | **81** | **114** | **485** |
