## Supplementary Table S3 for "Massive analysis of 64’628 bacterial genomes to decipher a water reservoir and origin of mobile colistin resistance (*mcr*) gene variants: is there another role for this family of enzymes?"

**Suppl. Table S3:** General features of depicted genes on the different genetic environment figures.

| | Gene name | Length (bp) | %GC content | Function |
|---|---|---|---|---|
| MCR-2 | DUF3987 | 1434 | 42.82 | DUF3987 domain-containing protein |
| | HP | 648 | 39.04 | hypothetical protein |
| | MCR-2 | 1617 | 47.19 | putative phosphatidylethanolamine transferase Mcr-1 |
| | IS630 | 1032 | 51.36 | IS630-like element ISSpu2 family transposase |
| | HP | 648 | 39.04 | hypothetical protein |
| | inovirus Gp2 | 534 | 45.32 | inovirus Gp2 family protein |
| | IS3 | 906 | 54.86 | IS3 family transposase |
| | Ion_trans2 | 408 | 39.95 | two pore domain potassium channel family protein |
| | PAP2 | 705 | 46.10 | PAP2 family lipid A phosphatase |
| | mcr1 | 1617 | 46.94 | putative phosphatidylethanolamine transferase Mcr-1 |
| | alpha/beta hydrolase | 1563 | 48.62 | alpha/beta hydrolase |
| | recombinase | 2268 | 46.34 | Recombinase |
| | acyl-CoA | 1776 | 50.17 | acyl-CoA dehydrogenase |
| MCR-3 | TnpA | 651 | 55.30 | TnpA transposase |
| | IS3 | 663 | 58.22 | IS3 family transposase |
| | Transposase | 315 | 56.19 | Transposase |
| | MCR-3 | 1626 | 41.76 | Phosphoethanolamine transferase EptA |
| | TnpA | 795 | 60.63 | TnpA transposase |
| | dgkA | 381 | 45.14 | Diacylglycerol kinase |
| | HP | 261 | 40.61 | hypothetical protein |
| | IS1 | 294 | 52.38 | IS1 family transposase |
| | HP | 216 | 54.63 | hypothetical protein |
| | IS6 | 132 | 48.48 | IS6 family transposase |
| | bleomycin binding | 318 | 41.19 | Bleomycin resistance protein |
| | HP | 261 | 40.61 | hypothetical protein |

| | | | | |
|---|---|---|---|---|
| MCR-4 | antitoxin | 288 | 38.19 | type II toxin-antitoxin system Phd/YefM family antitoxin |
| | antitoxin | 249 | 40.56 | type II toxin-antitoxin system Phd/YefM family antitoxin |
| | Hypothetical | 432 | 41.90 | hypothetical protein |
| | bin3 | 654 | 44.04 | Putative transposon Tn552 DNA-invertase bin3 |
| | Hypothetical | 210 | 38.10 | hypothetical protein |
| | HNH endonuclease | 309 | 33.01 | HNH endonuclease |
| | Tn3 transposase | 3087 | 41.43 | Tn3 transposase |
| | antitoxin | 312 | 43.27 | type II toxin-antitoxin system Phd/YefM family antitoxin |
| | antitoxin | 261 | 40.23 | type II toxin-antitoxin system Phd/YefM family antitoxin |
| | recombinase | 222 | 45.95 | recombinase |
| | recombinase | 279 | 41.94 | recombinase |
| | MCR-4 | 1626 | 40.10 | Phosphoethanolamine transferase EptA |
| | HNH endonuclease | 651 | 38.40 | HNH endonuclease |
| | Hypothetical | 204 | 37.75 | hypothetical protein |
| | Hypothetical | 291 | 37.46 | hypothetical protein |
| | Hypothetical | 237 | 37.97 | hypothetical protein |
| | Cobyrinic acid | 630 | 40.00 | Cobyrinic acid |
| MCR-5 | MFS | 657 | 64.99 | MFS transporter |
| | IS5 | 963 | 67.81 | IS5 family transposase |
| | MFS | 624 | 64.74 | MFS transporter |
| | MCR-5 | 1644 | 55.47 | Phosphoethanolamine transferase EptA |
| | chrB | 543 | 56.72 | Protein ChrB |
| | hin | 561 | 61.85 | DNA-invertase hin |
| | Tn3 transposase | 2967 | 63.67 | Tn3 transposase |
| MCR-8 | tnpA / IS5 | 924 | 52.71 | IS5 transposase family |
| | hhA | 204 | 38.73 | hemolysin expression modulator Hha |
| | thiJ | 699 | 47.78 | thiamine biosynthesis protein ThiJ |
| | GNAT | 510 | 53.14 | Acetyltransferase (GNAT) family protein |
| | GT | 912 | 37.06 | Glycosyl transferase |

| | | | | |
|---|---|---|---|---|
| | mcr-8 | 1698 | 40.40 | phosphoethanolamine--lipid A transferase MCR-8.1 |
| | copR | 696 | 47.84 | Transcriptional activator protein CopR |
| | baeS | 1230 | 43.01 | Integral membrane sensor signal transduction histidine kinase |
| | dgkA | 390 | 39.23 | Diacylglycerol kinase |
| | GT | 900 | 41.89 | Glycosyl transferase |
| | - | 210 | 45.24 | Transcriptional regulator |
| | - | 534 | 48.50 | hypothetical protein |
| | - | 357 | 42.58 | hypothetical protein |
| | ampC | 1092 | 54.85 | Beta-lactamase precursor |
| | - | 807 | 52.66 | MltA-interacting protein MipA |
| | sbmC | 471 | 46.28 | DNA gyrase inhibitor |
| | - | 90 | 40.00 | hypothetical protein |
| | ampC | 1365 | 48.50 | Beta-lactamase precursor |
| | tnpA / IS5 | 924 | 53.57 | IS5 transposase family |
| MCR-9 | *ATP/GTP* | 1206 | 47.51 | ATP/GTP-binding protein |
| | *DUF* | 819 | 49.21 | DUF4942 domain-containing protein |
| | *HP* | 168 | 42.26 | Hypothetical protein |
| | *rcnR* | 273 | 43.96 | Ni(II)/Co(II)-binding transcriptional repressor RcnR |
| | *rcnA* | 1116 | 49.73 | Nickel/cobalt efflux protein RcnA |
| | *pcoE* | 435 | 47.59 | Putative copper-binding protein PcoE |
| | *cusS* | 1347 | 47.14 | Sensor kinase CusS |
| | *IS5* | 924 | 54.11 | IS5 family transposase |
| | *mcr-9* | 1620 | 44.88 | Phosphoethanolamine transferase EptA |
| | *wbuC* | 477 | 53.04 | Cupin fold metalloprotein, WbuC family |
| | *hAMP* | 1350 | 50.37 | HAMP domain-containing histidine kinase |
| | *Res* | 669 | 53.21 | Response regulator |
| | *ATPAse* | 969 | 53.77 | AAA family ATPase |
| | *IS481* | 885 | 55.03 | IS481 family transposase |
| | *IS6* | 705 | 53.33 | IS6 family transposase |
| | *IS110* | 1023 | 49.76 | IS110 family transposase |
| | *toxin-antitoxin* | 330 | 50.00 | Type II toxin-antitoxin system RelE/ParE family toxin |

| | | | | |
|---|---|---|---|---|
| | *ardK* | 342 | 43.57 | Transcriptional regulator ArdK |
| | *zinc M* | 792 | 52.02 | Zinc metalloprotease |
| | *aph(6)-I* | 837 | 55.91 | APH(6)-I family aminoglycoside O-phosphotransferase |
| | *aph(3'')-Ib* | 804 | 56.22 | Aminoglycoside O-phosphotransferase APH(3'')-Ib |
| | *xerD* | 1014 | 61.14 | Tyrosine recombinase XerD |
| | *endonuclease* | 642 | 49.07 | Restriction endonuclease |
